# BRAPH: A Graph Theory Software for the Analysis of Brain Connectivity

**DOI:** 10.1101/106625

**Authors:** Mite Mijalkov, Ehsan Kakaei, Joana B. Pereira, Eric Westman, Giovanni Volpe, for the Alzheimer’s Disease Neuroimaging Initiative

## Abstract

The brain is a large-scale complex network whose workings rely on the interaction between its various regions. In the past few years, the organization of the human brain network has been studied extensively using concepts from graph theory, where the brain is represented as a set of nodes connected by edges. This representation of the brain as a connectome can be used to assess important measures that reflect its topological architecture. We have developed a freeware MatLab-based software (BRAPH – BRain Analysis using graPH theory) for connectivity analysis of brain networks derived from structural magnetic resonance imaging (MRI), functional MRI (fMRI), positron emission tomography (PET) and electroencephalogram (EEG) data. BRAPH allows building connectivity matrices, calculating global and local network measures, performing non-parametric permutations for group comparisons, assessing the modules in the network, and comparing the results to random networks. By contrast to other toolboxes, it allows performing longitudinal comparisons of the same patients across different points in time. Furthermore, even though a user-friendly interface is provided, the architecture of the program is modular (object-oriented) so that it can be easily expanded and customized. To demonstrate the abilities of BRAPH, we performed structural and functional graph theory analyses in two separate studies. In the first study, using MRI data, we assessed the differences in global and nodal network topology in healthy controls, patients with amnestic mild cognitive impairment, and patients with Alzheimer’s disease. In the second study, using resting-state fMRI data, we compared healthy controls and Parkinson’s patients with mild cognitive impairment.

## 1. Introduction

Graph theory studies the properties and behavior of networks, which are systems consisting of a set of elements (nodes) linked by connections or interactions (edges). Many systems found in Nature, ranging from social interactions to metabolic networks and transportation systems, can be modeled within this framework, pointing to a set of underlying similarities among these very diverse systems. The human brain can also be modeled as a network (the *human connectome*) [1], where brain regions are the *nodes* and the connections between them are the *edges*. The human brain is thus an ideal candidate for graph theoretical analysis. The nodes can be defined as the brain regions underlying electrodes or using an anatomical, functional or histological parcellation scheme. The edges are obtained as measures of association between the brain regions, such as connection probabilities (diffusion tensor imaging, DTI), inter-regional correlations in cortical thickness (magnetic resonance imaging, MRI) and electrophysiological signals (electroencephalography, EEG; magnetoencephalography, MEG) or statistical dependencies in time series (functional MRI, fMRI) and blood flow (arterial spin labeling, ASL).

After compiling all pairwise associations between the nodes into a *connectivity matrix* (or *brain graph*), several network properties can be calculated in order to characterize the global and local organization of the connectome. For instance, the *small-worldness* can be used to assess the balance between short-distance and long-distance connectivity [2], while the *modularity* defines how well the network can be divided into *subnetworks* (or *modules*) [3,4], which generally correspond to well-known brain systems such as the default-mode or fronto-parietal networks [5,6]. These network properties and many others can be used to reveal fundamental aspects of normal brain organization and highlight important aspects of underlying brain pathology in diseases such as Alzheimer’s disease (AD) [7] or Parkinson’s disease (PD) [8].

Several toolboxes have been developed to study brain connectivity, including the Brain Connectivity Toolbox [9], eConnectome [10], GAT [11], CONN [12], BrainNet Viewer [13], GraphVar [14] and GRETNA [15]. While all of them made important contributions by proving new options to build, characterize and visualize brain network topology, they require some programming experience, or deal only with some aspects of the analysis, or are coded in such a way that their adaptation is hard to achieve. Hence, a reliable, streamlined, user-friendly, fast, and scalable software that deals with all aspects of network organization is still lacking.

In this article, we present BRAPH – BRain Analysis using graPH theory (http://www.braph.org/), a software package to perform graph theory analysis of the brain connectome. BRAPH is the first object-oriented open-source software written in MatLab for graph theoretical analysis with a graphical user interface (GUI). In contrast to previous toolboxes, BRAPH takes advantage of the object-oriented programming paradigm to provide a clear modular structure that makes it easy to maintain and modify existing code, since new objects can be added without the need for an extensive knowledge of the underlying implementation. From the clinical point of view, BRAPH presents the following strengths: (a) it allows comparing regional node values before the actual network analysis, which is important to get a first impression on the data; (b) it visualizes individual connectivity matrices and individual network measures, which is crucial to detect potential outliers, a major confound in neuroimaging studies; (c) it carries out longitudinal graph theory analyses that provide an important insight into topological network changes over time; (d) it assesses modular structure using different algorithms and allows performing subnetwork analyses within the defined modules, which is important for studies testing hypotheses within a particular structural or functional brain network; (e) it provides utilities for multi-modal graph theory analyses by integrating information from different network modalities, which is arguably the next challenge in imaging connectomics. From the user point of view, BRAPH is the only fully vertically integrated software that allows carrying out all the steps of a graph theory analysis, from importing the neuroimaging data to saving the final results in a single file. Importantly, this is not only practical but also increases the reliability and reproducibility of the results, which is an increasingly important issue within the research community. In addition, BRAPH offers a comprehensive manual, continuous release of new utilities, and a support forum, all of which can be accessed at http://www.braph.org/. Finally, BRAPH offers online videos that provide a step-by-step guide on how to perform graph theory analyses, allowing the users a simple and quick start for their brain connectivity studies.

Below we describe in detail the different options that BRAPH offers for graph theory analyses when it comes to building connectivity matrices, applying threshold strategies, performing weighted or binary network analyses and computing random networks. Our software has already been successfully applied in previous graph theory studies [16,17] but to further demonstrate its abilities, in this article we assess network topology on structural MRI data from patients with amnestic mild cognitive impairment (MCI) and AD, and on fMRI data from PD patients with MCI.

## 2. Materials and Methods

### 2.1. Overview of BRAPH

BRAPH is a complete software package that allows carrying out all the steps of a graph theoretical analysis, visualize the results and generate high-quality publication-ready images. It can obtain undirected binary and weighted brain connectivity graphs starting from data acquired using various neuroimaging modalities, including MRI, fMRI, EEG, and positron emission tomography (PET). BRAPH can also assess the modular structure of the brain graph, employing various algorithms and extracting modules for further analysis. To test for significant differences between groups, BRAPH carries out non-parametric permutation tests and allows correcting the results for multiple comparisons using false discovery rate (FDR) [18]. It also provides options to carry out longitudinal graph theory analyses and to normalize the network measures by random graphs.

As shown in Figure 1, the software consists of three independent layers connected by software interfaces: Graph, Data Structures and Graphical User Interfaces (GUIs). The Graph package includes the fundamental functions to perform a graph theory analysis and calculating the global and nodal measures. The Data Structures package provides the core functionalities of the software and allows defining the brain atlas, the cohort of subjects, and the type of graph analysis; importantly, all these functionalities can be accessed by command-line and can therefore be scripted by advanced users. Finally, the GUIs package provides a streamlined way to carry out graph theory analyses based on a series of GUIs for users without a computational background: (a) the *GUI Brain Atlas* allows selecting and editing the brain atlas; (b) the *GUI Cohort* allows defining the cohort of subjects by uploading the relevant data; (c) the *GUI Graph Analysis* allows building the connectivity matrices by selecting the type of graph (*weighted*, *binary*, see also Figure 2) and thresholding method (*threshold*, *density*) as well as calculating topological measures and visualizing the results. For the *GUI Cohort* and *GUI Graph Analysis*, four options can be selected (MRI, fMRI, PET, EEG) depending on the nature of the analysis. Figure 3 shows an overview of these different steps. Thanks to this three-layered structure, BRAPH can be easily expanded and customized to address any needs, e.g., by implementing new graph measures or new approaches for building brain graphs.

**Figure 1.**
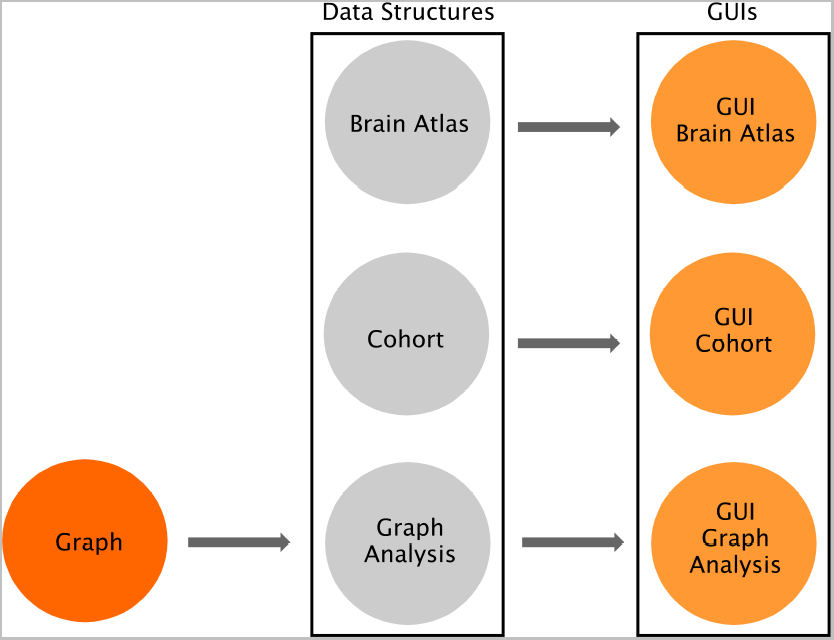
Overview of BRAPH software architecture. BRAPH consists of three layers, from left to right: *Graph*, *Data Structures* and *Graphical User Interfaces* (GUIs). These layers are connected by unidirectional software interfaces (arrows). *Graph* contains the functions to perform graph analyses. In *Data Structures*, *Brain Atlas* allows defining the nodes of the network, *Cohort* allows defining the subjects to be studied and dividing them into groups, and *Graph Analysis* permits building the connectivity matrices and calculating network measures; each of these is implemented in an object, whose functionalities can be called by command line. For each of these objects, a GUI is provided (i.e. *GUI Brain Atlas*, *GUI Cohort* and *GUI Graph Analysis*). Thanks to this architecture BRAPH can be very easily maintained, expanded and customized.

**Figure 2.**
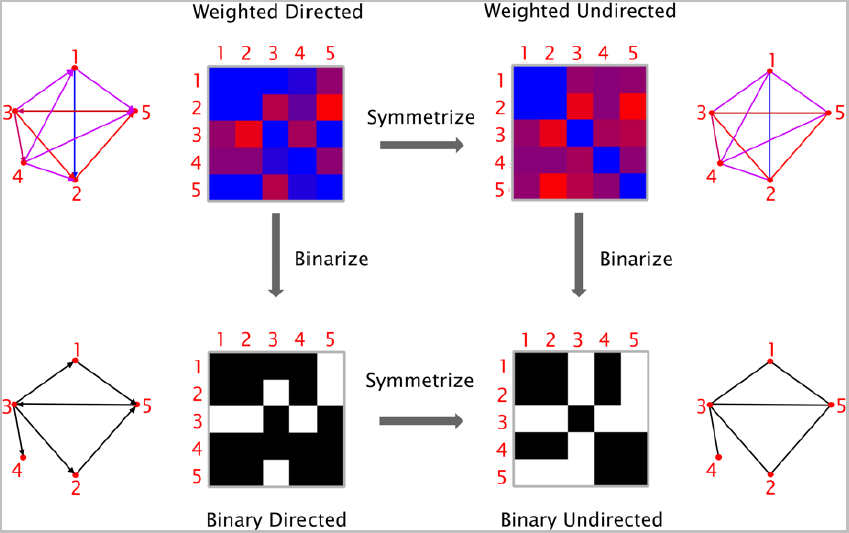
Types of graphs. Graphs can be classified based on their edge weights (*weighted* or *binary*) and directionality (*directed* or *undirected*). It is possible to transform a directed graph into an undirected one by symmetrization (i.e. by removing the information about the edge directions), and a weighted graph into a binary one by thresholding (i.e. by assigning a value of 1 to the edges above a given threshold and 0 to those below threshold).

**Figure 3.**
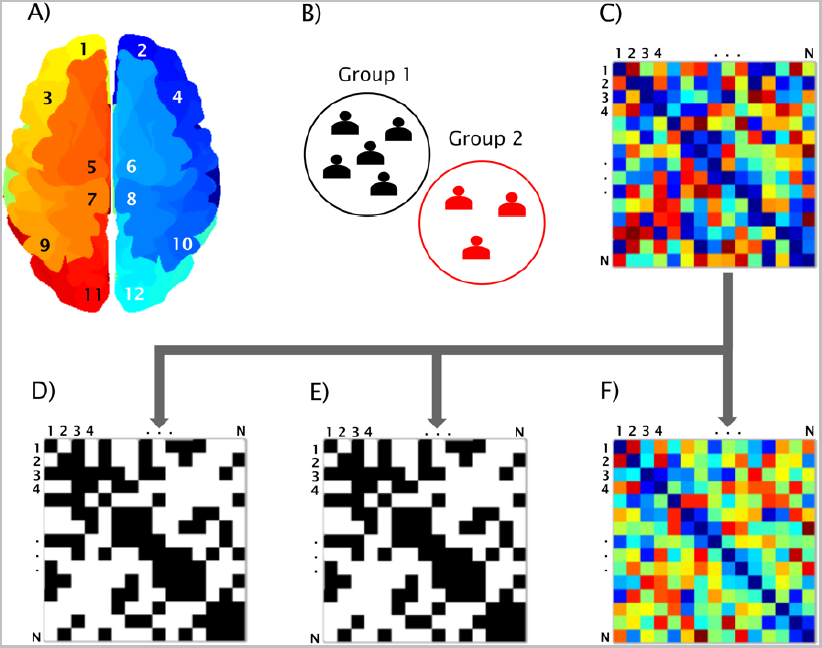
BRAPH workflow. Workflow for a graph theory analysis in BRAPH and relative graphical user interfaces (GUIs). A) The brain regions are defined in the *GUI Brain Atlas*. B) The data of the subjects are imported in the *GUI Cohort* and the user can define groups and edit their age, gender and other relevant data. C) The connectivity matrix is calculated in the *GUI Graph Analysis* after selecting the parameters defining the type of correlation, how to deal with negative correlation coefficients, and which type of graph to analyze: D) binary undirected graphs at a fixed density (*GUI Graph Analysis BUD*); E) binary undirected graphs at a fixed threshold (*GUI Graph Analysis BUT*); F) weighted undirected graphs (*GUI Graph Analysis WU*).

### 2.2. Defining the nodes

The first step of a graph theory analysis consists of defining the nodes, which generally correspond to the regions included in a brain atlas. This atlas should contain the names and labels of the brain regions as well as their spatial coordinates (*x*, *y*, *z*) in order to project them on a 3D surface and create a visual representation of the brain graph. In the case of the analysis of structural networks (e.g. obtained from structural MRI data or T1-weighted images), the nodes are usually defined using an anatomical parcellation scheme that divides the brain into regions using the brain sulci and gyri as anatomical landmarks. Examples of anatomical atlases are the automated anatomical labeling (AAL) [19], Desikan [20] or Destrieux [21] atlases. BRAPH already provides these atlases ready for upload on the *GUI Brain Atlas* interface. In the case of the analysis of functional networks (e.g. obtained from fMRI data), the atlas may be defined using an anatomical parcellation scheme, a meta-analysis, or a clustering-based method of spatially coherent and homogenous regions. Examples of functional atlases are the Dosenbach [22], the Power [23] and the Craddock [24] atlases, all of which are also provided by BRAPH. The user may also upload a different atlas from an external file (in .xml, .txt or .xls format) or create an entirely new one in the GUI. The resulting atlas can be saved as a .atlas file (see manual and website for an example of a .atlas file).

After the atlas has been created or uploaded into the software, the user should then upload the subject data into the *GUI Cohort* interface. This data may consist, for example, of cortical thickness, surface area or volume measures in structural MRI; regional time-series in resting-state fMRI; glucose metabolism or blood flow in PET and ASL; and electrophysiological signals in EEG or MEG. These regional values can be obtained using the Statistical Parametric Mapping (SPM; http://www.fil.ion.ucl.ac.uk/spm/), FMRI Software Library (FSL; https://fsl.fmrib.ox.ac.uk/fsl/fslwiki), FreeSurfer (https://surfer.nmr.mgh.harvard.edu/) or any other image preprocessing software. In addition, they may be corrected for the effects of nuisance variables such as age, gender or scanner site by means, for example, of linear regression (in this case the residual values should substitute the raw values in the network analysis) [25].

### 2.3. Defining the edges

Once the nodes of the network have been defined, the edges representing the relationship between them need to be computed. In BRAPH, the edges are calculated in *GUI Graph Analysis* as the statistical correlation between the values of all pairs of brain regions for an individual or for a group of subjects, depending on the neuroimaging technique. Different types of parametric and non-parametric correlations may be selected for this purpose: Pearson, Spearman, Kendall rank correlation coefficients, or (Pearson or Spearman) partial correlation coefficients. Note that all self-connections are eliminated from the analysis by setting the diagonal entries in the connectivity matrix to zero. In addition, the user can also choose whether to retain the negative correlation coefficients, substitute them with their absolute value or replace them by zero.

### 2.3. Network construction

To minimize the computation time, the graph measures can be calculated using optimized algorithms based on linear algebra. Therefore, a graph is more conveniently represented as a connectivity matrix, where the rows/columns denote the nodes and the matrix elements represent the edges between the nodes. Each row of the connectivity matrix represents the edges that are going out from a node; for example, row *j* represents the edges that are going out from node *j*. Each column of the matrix represents the nodes that arrive to a node; for example, column *k* represents the edges that are arriving to node *k*. Thus, the element (*j*, *k*) represents the edge that goes from node j to node k. The specific order of the nodes in the matrix does not affect the calculation of the graph theory measures, but only the graphical representation of the connectivity matrix. As illustrated in Figure 2, based on the nature of the edge’s weight and directionality, four types of graphs can be defined. *Weighted directed* (WD) graphs have edges associated with a real number, indicating the strength of the connection, and are directed (i.e., node *j* can be connected to node *k* without node *k* being connected to node *j*). The edges in the *weighted undirected* (WU) graphs are associated with a real number indicating the strength of the connection and are undirected (i.e., if node *j* is connected to node *k*, then node *k* is also connected to node *j*), resulting in a symmetric connectivity matrix. *Binary directed* (BD) graphs have directed edges, which can either be 0 or 1, indicating the absence or presence of a connection. The edges in a *binary undirected* (BU) network can also be either 0 or 1 and they have no preferential directionality. In order to transform a directed graph into an undirected graph, the connectivity matrix needs to be symmetrized. In BRAPH, the connectivity matrix can be symmetrized via command line by: (a) taking the sum between the matrix itself and its transpose; (b) taking the difference between the previous two; (c) comparing the matrix to its transpose and selecting either the smaller or the larger value for each entry. We remark that, even though the directed measures are not currently used in the analyses performed by BRAPH, they are already available in the *Graph* package and ready to be used in future versions of the software.

To transform a weighted graph into a binary one, BRAPH assigns a value of 1 to the edges above a given threshold and 0 to those below it. There are two ways of applying a threshold: (a) by selecting a correlation coefficient as the cut-off value below which all connections are excluded from the analysis (*binary undirected threshold* (BUT) interfaces); or (b) by fixing the fraction of edges (i.e., a specific density) that will be connected (*binary undirected density* (BUD) interfaces). The choice between these two options becomes significant when comparing different groups of subjects, as it may lead to different results; currently the density approach is more often employed in the literature, because it permits analyzing differences in network architecture, while controlling for the different number of edges across individuals or groups.

For MRI or static PET data, a single connectivity matrix is calculated for each group of subjects; therefore, the graph theory measures reflect the group’s properties. For fMRI data and other neuroimaging sequences that provide a measure of brain function over time, an individual connectivity matrix is calculated for each subject; therefore, the graph theory measures reflect the characteristics of each subject, which can then be averaged within a particular group.

### 2.4. Network analysis

BRAPH allows calculating both global and nodal network measures, on weighted or binary networks, using different thresholds or densities. To test for significant differences between groups (cross-sectional analysis) or two different points across time (longitudinal analysis), BRAPH performs non-parametric permutation tests, reporting one-tailed and two-tailed p-values based on 95% confidence intervals.

The network measures can also be compared with the corresponding measures calculated on random graphs with the same degree or weight distribution. These can be used, for example, to normalize weighted network measures.

Regarding nodal network measures, the permutation tests are carried out for each brain region, assessing simultaneously multiple null hypotheses, which consequently increases the risk of finding false positives. BRAPH deals with this issue by providing the adjusted p-values that should be considered to correct the results for multiple comparisons with false discovery rate (FDR) using the Benjamini-Hochberg procedure [18].

### 2.5. Graph theory measures

BRAPH can calculate several graph theory measures that assess the topology of the whole brain network as well as of its regions. Here, we explain briefly some of the most relevant ones; for a complete list with formulas and details, please refer to the BRAPH manual and website. The code we used to calculate the graph measures was adapted from the Brain Connectivity Toolbox (http://www.brain-connectivity-toolbox.net/) [9], which is regarded as the most important reference in the field since it provided the seminal groundwork for the use of graph theory by neuroimaging researchers.

The simplest, yet most fundamental, measure that can be assessed in a graph is the *degree*, which is the number of connections a node has with the rest of the network. In weighted graphs, we calculate the degree of the nodes by ignoring the weights and binarizing the matrix. The degree distribution in the brain follows a power law [26], meaning that highly connected areas tend to communicate with each other.

Another important measure is the *shortest path length*, which is the shortest distance between two nodes. In a binary graph, distance is measured as the minimum number of edges that need to be crossed to go from one node to the other. In a weighted graph, the length of an edge is a function of its weight; typically, the edge length is inversely proportional to the edge weight because a high weight implies a stronger connection. The average of the minimum path lengths between one node and all other nodes is the *characteristic path length* [2]. One can also define two related measures of centrality: the *closeness centrality*, which is the inverse of the shortest path length, and the *betweenness centrality*, which is the fraction of all shortest paths in the network that pass through a given node [9]. These and other measures can be used to assess whether a node is a brain hub [27], regulating most of the information flow within the network.

The closer the nodes are to each other, the shorter is the path length and the more efficient is the transfer of information between them. Therefore, one can define the *global efficiency* of a node as the inverse of the shortest path from that node to any other node in the network [28]. To assess the communication efficiency between a node and its immediate neighbors, the *local efficiency* can be calculated. Both global and local efficiency measures can be averaged over all nodes to describe global properties of the brain network [28].

The *clustering coefficient* is a measure that assesses the presence of cliques or clusters in a graph [2]. For each node, this can be calculated as the fraction of the node’s neighbors that are also neighbors of each other. For the whole network, the nodal clustering coefficients of all nodes can be averaged into the mean clustering coefficient. A closely related measure to the clustering is the *transitivity*, which is defined as the ratio of paths that transverse two edges by the number of triangles. If a node is connected to another node, which in turn is connected to a third one, the transitivity reflects the probability that the first node is connected to the third.

The *small-worldness* is given by the ratio between the characteristic path length and mean clustering coefficient (normalized by the corresponding values calculated on random graphs) [2]; this is an important organizational property that describes an optimal network architecture. Compared to a random graph, a small-world network is characterized by similarly short paths but a significantly higher clustering coefficient.

A network can also be divided into separate communities corresponding to anatomical proximity or to a specific function shared by a group of nodes. The extent to which a network can be divided into these communities or modules can be calculated using the *modularity*, which maximizes the number of edges within communities and minimizes the number of edges between different communities [29]. The *within-module z-score* quantifies how well a node is connected with other nodes from the same module, while the *participation coefficient* assesses if a node has many connections with nodes from different modules. If a node has a high within-module degree it is classified as a provincial hub; if it has a high participation coefficient it is considered to be a connector hub.

### 2.6. Subjects

To demonstrate the abilities of BRAPH, we performed structural and functional graph theory analyses in two separate studies. In the first study, we assessed the differences in global and nodal network topology in healthy controls, patients with amnestic MCI, and patients with AD (see Table 1) from the Alzheimer’s Disease Neuroimaging Initiative (ADNI) database (adni.loni.usc.edu). The ADNI was launched in 2003 as a public–private partnership, led by Principal Investigator Michael W. Weiner, MD. The primary goal of ADNI has been to test whether serial MRI, PET, other biological markers, and clinical and neuropsychological assessment can be combined to measure the progression of MCI and early AD. All participants were scanned on a 1.5 Tesla MRI system using a sagittal 3D *T*_1_-weighted MPRAGE sequence: repetition time (TR) = 9–13 ms; echo time (TE) = 3.0–4.1 ms; inversion time (IT) = 1000 ms; flip angle (FA) = 8°; voxel size = 1.1 × 1.1 × 1.2 mm3.

**Table 1.** Characteristics of the structural MRI sample.

| | CTR<br>(n=42) | MCI<br>(n=42) | AD<br>(n=48) | F or $\chi^2$<br>(p value) |
| --- | --- | --- | --- | --- |
| Age (y) | 76.1(5.0) | 74.5(7.5) | 75.6 (7.0) | 0.017 |
| Sex (M/F) | 110/100 | 243/134 | 97/84 | 0.005 |
| Education (y) | 16.0(2.9) | 15.7(3.0) | 14.8(3.2) | <0.001 |
| MMSE | 29.1(0.9) | 27.0(1.8) | 23.2(2.0) | <0.001 |

Means are followed by standard deviations. Differences in age, years of education, and MMSE scores were assessed using an analysis of variance (ANOVA). Differences in gender were assessed using a $\chi^2$ test. CTR, controls; MCI, mild cognitive impairment; AD, Alzheimer's disease; MMSE, mini-mental state examination.

In the second study, we carried out a graph theory analysis on the resting-state fMRI data of healthy controls and PD patients with MCI (see Table 2) from the Parkinson’s Progression Markers Initiative (PPMI) (2011) [30] (http://www.ppmi-info.org/data; accessed in November, 2015), an international, multicenter study launched in 2010 to identify PD progression biomarkers. Each participating PPMI site received approval from an ethical standards committee before study initiation and obtained written informed consent from all participants. PD patients were classified as having MCI (PD-MCI) if they scored 1.5 standard deviations below the scaled mean scores on any two cognitive tests, following previously published procedures for PD-MCI diagnosis in the PPMI cohort [31] PD-MCI patients and controls were scanned on a 3 Tesla Siemens scanner (Erlangen, Germany). Resting-state functional images were acquired using an echo-planar imaging sequence (repetition time = 2400 ms; echo time = 25 ms; flip angle = 80⁰; matrix = 68 x 68; voxel size = 3.25 x 3.25 x 3.25 mm3). The scan lasted 8 minutes and 29 seconds and included 210 volumes.

**Table 2.** Characteristics of the fMRI sample.

|  | CTR<br>(n=15) | PD-MCI<br>(n=15) | CTR vs PD-MCI<br>(p value) |
| --- | --- | --- | --- |
| Age (y) | 66.4(9.1) | 63.5(8.2) | 0.372 |
| Sex (M/F) | 13/2 | 11/4 | 0.361 |
| Education (y) | 16.5(2.3) | 14.2(3.1) | 0.024 |
| UPDRS-III | - | 21.8(8.9) | - |
| HY stage | - | 1.8(0.4) | - |
| MoCA | 24.0(3.7) | 27.9(1.6) | 0.001 |

Means are followed by standard deviations. Differences in age, years of education, and MoCA scores were assessed using Student's T test. Differences in gender were assessed using a $\chi^2$ test. CTR, controls; PD-MCI, Parkinson's disease with mild cognitive impairment; UPDRS-III, Unified Parkinson's disease rating scale - Part III; HY stage, Hoehn and Yahr stage; MoCA, Montreal cognitive assessment scale.

### 2.7. Network construction and analysis

To assess structural network topology in controls, amnestic MCI patients, and AD patients from ADNI, the T1-weighted images of these subjects were preprocessed using FreeSurfer (version 5.3), as published elsewhere [17]. The cortical thickness and subcortical volumes of 82 regions were extracted and included as nodes in the network analysis. The edges between these regions were computed as Pearson correlations and the network analyses were carried out on the binary undirected graphs, while controlling for the number of connections, across a range of densities: from 5% to 25%, in steps of 0.5%.

To assess functional network topology in PD-MCI patients and a group of elderly controls from PPMI, fMRI images were preprocessed using SPM8 (http://www.fil.ion.ucl.ac.uk/spm) using the following steps: removal of first five volumes, slice-timing correction, realignment, normalization to the Montreal Neurological Institute (MNI) template (voxel size 3x3x3mm^3^), temporal filtering (0.01-0.08 Hz), regression of white matter, cerebrospinal fluid signals and six head motion parameters. The regional time-series of the 200 brain regions included in the Craddock atlas [24] were extracted from each subject. To compute the relationship between these regions, we used Pearson correlations and performed the network analyses on the weighted undirected graphs.

In both studies, non-parametric permutation tests were carried out to assess differences between groups, which were considered significant for a two-tailed test of the null hypothesis at *p*<0.05. In addition, to adjust the nodal network results for multiple comparisons, a FDR procedure was applied to control for the number of regions that were tested at *q*<0.05.

## 3. Results

### 3.1. Structural network topology in amnestic MCI and AD

The structural correlation matrices and brain graphs of patients and controls can be found in Figure 4. All groups showed strong correlations between bilaterally homologous regions.

**Figure 4.**
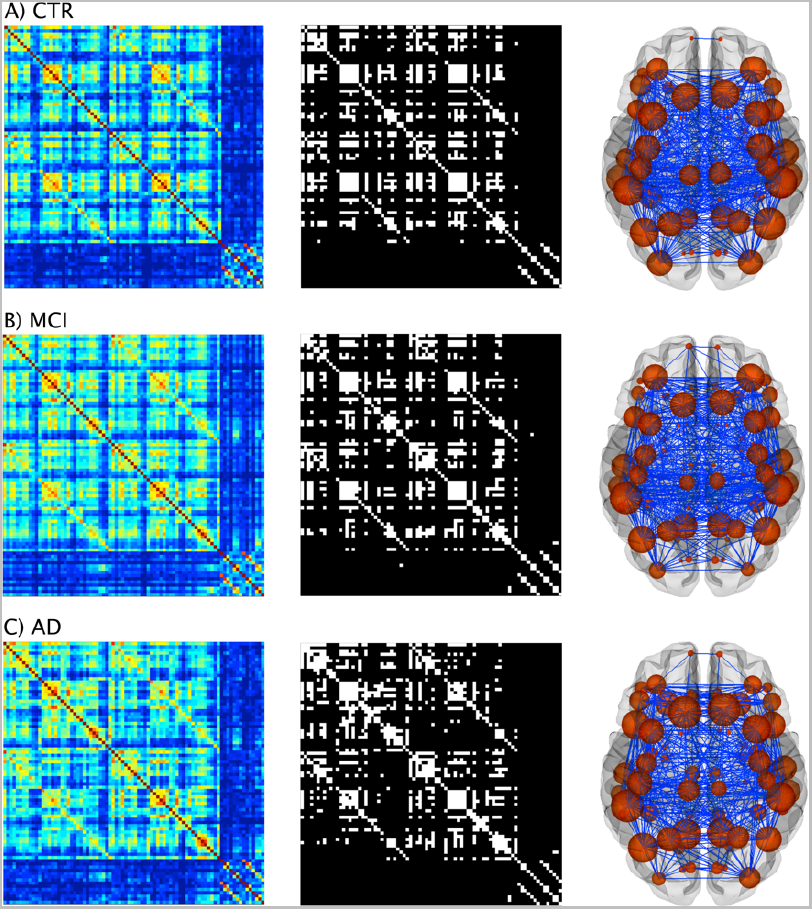
Structural brain networks in controls, MCI patients, and AD patients. From left to right: weighted correlation matrices of 82 regions, binary correlation matrices after fixing density at 15%, and corresponding brain graphs from A) controls (CTR), B) patients with amnestic mild cognitive impairment (MCI), and C) Alzheimer’s disease (AD) patients.

Regarding global network topology (Figure 5), we found increases in the characteristic path length and local efficiency in MCI and AD patients compared to controls at several network densities. The transitivity and modularity showed the most widespread topological changes: the transitivity was decreased and the modularity was increased in MCI and AD patients across almost all network densities compared to controls. Compared to MCI, AD patients showed increases in the characteristic path length at a few network densities and widespread changes in the transitivity and modularity.

**Figure 5.**
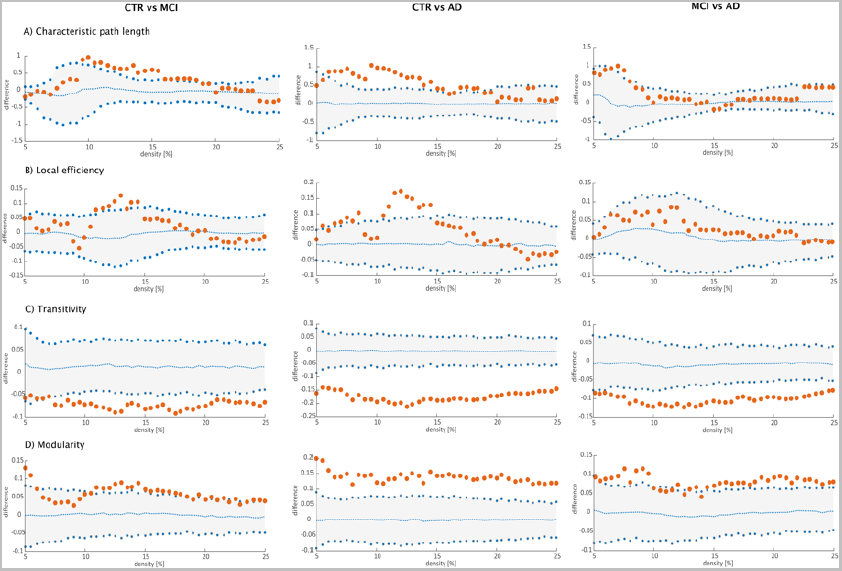
Differences between groups in global structural topology. Left: differences between controls (CTR) and Alzheimer’s disease (AD) patients; middle: differences between controls (CTR) and patients with mild cognitive impairment (MCI); right: differences between patients with mild cognitive impairment (MCI) and Alzheimer’s disease (AD) patients for A) characteristic path length, B) local efficiency, C) transitivity and D) modularity. The plots show the lower and upper bounds (blue circles) of the 95% Confidence Intervals (CI) (gray shade) as a function of density. The orange circles show the differences between groups and, when falling outside the CI, indicate that the difference was statistically significant at *p*<0.05. The blue dots indicate the mean values of the difference in global network measures between the randomized groups after permutation tests.

**Figure 6.**
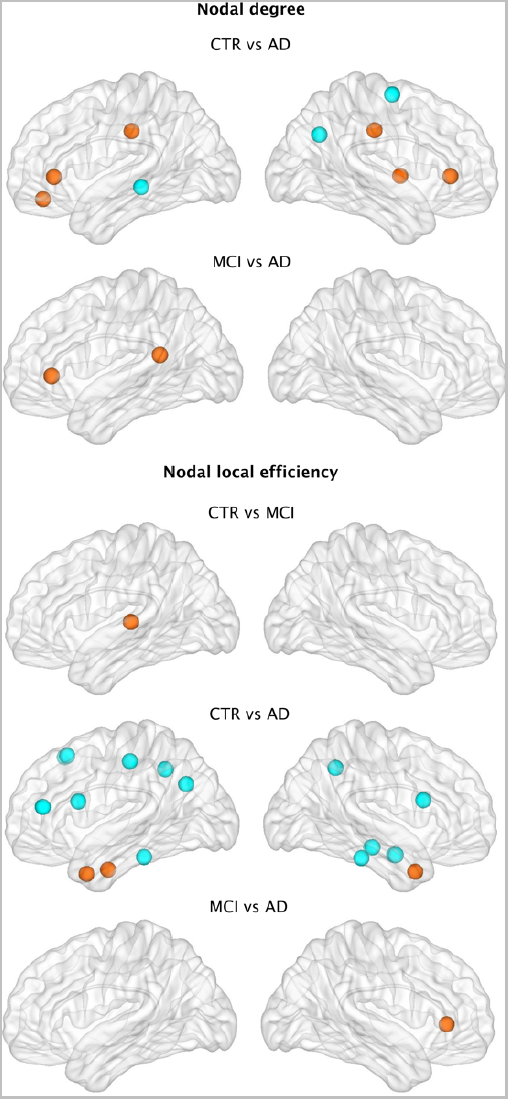
Differences between groups in nodal structural measures. Nodes showing significant differences between groups in the nodal degree and nodal local efficiency after FDR corrections. Orange indicates increases in the nodal measure, while blue indicates decreases.

Regarding regional network topology (Figure 6), the nodal degree showed significant increases in the left medial orbitofrontal, right insula, bilateral rostral anterior cingulate and posterior cingulate gyri in addition to decreases in the left middle temporal, right precentral and right inferior parietal gyri in AD patients compared to controls. When compared to MCI, AD patients also presented a higher nodal degree in the left rostral anterior cingulate and isthmus cingulate gyri.

We also compared the nodal local efficiency between groups. This measure showed significant increases in the left transverse temporal gyrus in MCI patients compared to controls. AD patients showed both increases in the local efficiency in the bilateral temporal pole and left entorhinal cortex as well as decreases in several regions from the frontal (bilateral superior frontal, left pars triangularis, bilateral pars opercularis, right postcentral gyri), temporal (bilateral inferior temporal gyri, amygdala, hippocampus) and parietal (left inferior parietal, right precuneus) lobes. When the two patient groups were compared to each other, AD patients showed efficiency increases in the right rostral anterior cingulate compared to the MCI group.

### 3.2. Functional network topology in PD-MCI

The functional connectivity matrices of controls and PD-MCI patients can be found in Figure 7. The comparison of the weighted average degree showed that PD-MCI patients presented a significantly lower number of connections compared to controls (PD-MCI=172.6; controls=183.7; p-value=0.027), suggesting that their network was more disconnected. Then, we performed a modularity analysis on the weighted graphs of these subjects to assess the presence of smaller communities of regions (modules). This analysis showed there were five modules in controls and patients, which were quite similar in the two groups. Module I included medial frontal areas, the posterior cingulate and bilateral angular gyri, resembling the default-mode network. Module II comprised temporal and cerebellar areas. Module III included several middle, inferior frontal and parietal regions, similarly to the fronto-parietal network usually found in resting-state studies [5,6]. Module IV consisted of most of the visual cortex similarly to the previously reported visual network. Finally, Module V included mainly subcortical regions and a few temporal and cingular areas.

**Figure 7.**
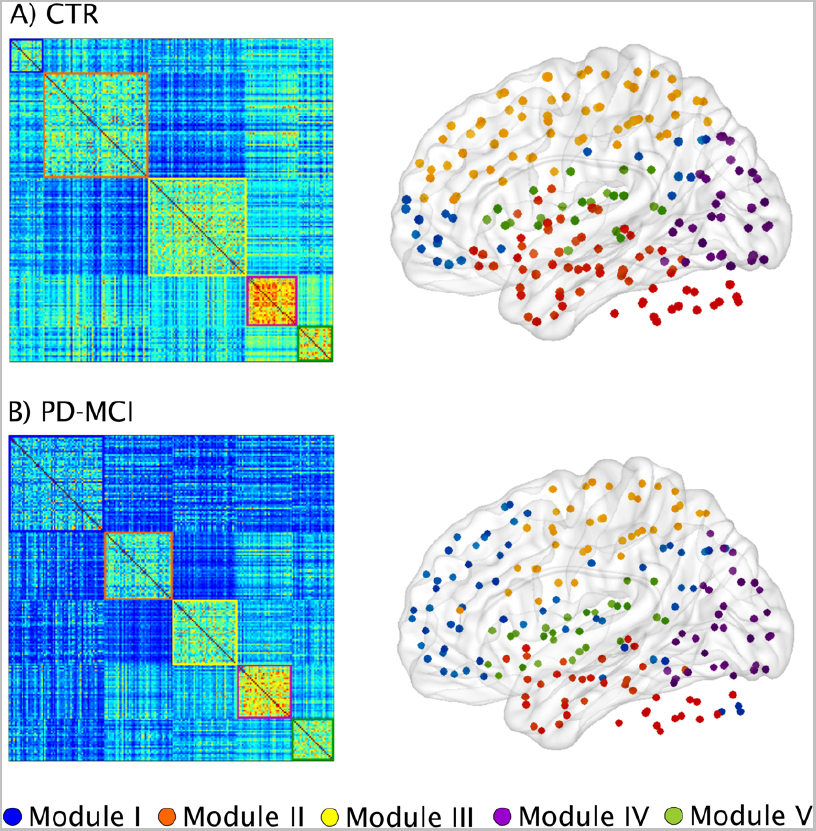
Functional brain networks and modules in controls and PD-MCI patients. Weighted connectivity matrices and modules in A) controls (CTR) and B) Parkinson’s disease patients with mild cognitive impairment (PD-MCI). Five modules were identified in each group.

When we assessed the nodal degrees of the regions included in each module (subgraph analysis, Figure 8), we observed that the only differences between the control and PD-MCI groups were found in the fronto-parietal network. Within this network, the regions showing a significantly lower degree in PD-MCI patients after FDR corrections were the bilateral superior frontal, superior parietal gyri and precuneus in addition to the left middle and inferior frontal gyri and anterior cingulate.

**Figure 8.**
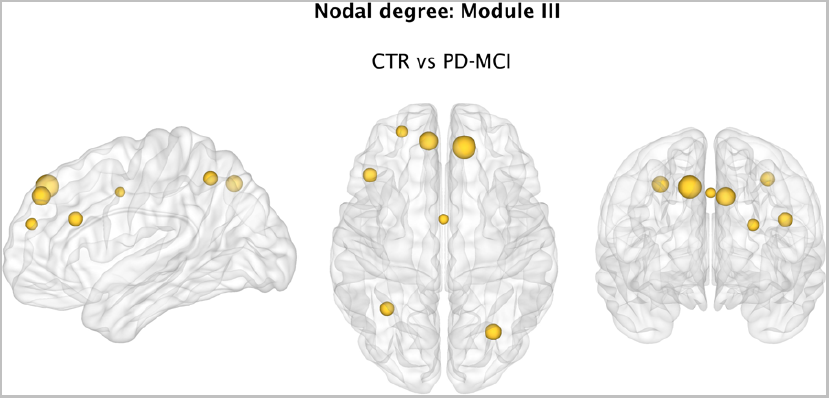
Differences between groups in the nodal functional degree. Significant decreases in the nodal degree of regions from Module III or fronto parietal network in Parkinson’s disease patients with mild cognitive impairment (PD-MCI) compared to controls (CTR) after FDR corrections.

## 4. Discussion

Graph theory has introduced new opportunities for understanding the brain as a complex system of interacting elements. Thanks to this framework, we have come to appreciate that the human brain relies on fundamental aspects of network organization such as a small-world architecture, modular structure and vulnerable hubs. These properties allow our brains to evolve, grow and adapt within an environment presenting increasing cognitive demands, and their disruption accounts for some of the key aspects underlying pathology in neurological diseases. In this report, we present BRAPH, the first object-oriented software for graph theory analysis intended for all researchers, regardless of their scientific background. As modern network science continues to develop at an amazing speed, it is important to have a software that allows modifying existent code in a structured manner so that past knowledge can be easily integrated with new topological analyses and graph theory measures. We are currently working on some of these developments with a particular focus on multimodal analyses and effective connectivity measures. Amongst BRAPH’s strengths is the fact that it deals with all the aspects of graph theory analysis, by providing the user with extensive assistance, from the first basic steps such as defining the nodes or edges to producing the final end-stage figures of the results as well as to archiving the results and relative analysis procedure in a dedicated file. To get an impression on BRAPH’s abilities, below we discuss some of the results we obtained in two different studies in patients with amnestic MCI, AD and PD-MCI.

### 4.1. Large-scale structural networks in amnestic MCI and AD

AD is currently one of the most prevalent neurodegenerative disorders, with a significant impact on society and caregiver burden [32]. Although the devastating impact of this condition has pushed forward a large research effort towards a more accurate diagnosis, the underlying effects of AD on network topology remain poorly understood. There is increasing evidence suggesting that the pathological hallmarks of AD, consisting of amyloid plaques neurofibrillary tangles, could spread in the brain through synapses and neural connections. Hence, the application of graph theory to the study of brain connectivity could shed light on the mechanisms of disease propagation in AD.

In the current study, we found that AD patients presented an abnormal global network topology as reflected by increases in the path length, local efficiency and modularity, and by decreases of transitivity. These changes indicate that the regions of their networks communicated less efficiently with other brain regions and between different brain modules. In particular, the most widespread changes in network organization were observed for the transitivity and modularity. The decreases in transitivity found in AD suggest that the regions of their network were poorly connected to neighboring areas, whereas the increases in modularity indicate that their modules had higher within-module connectivity and worse inter-module connectivity. Patients with amnestic MCI, who are potentially on the path to develop AD, also showed similar, albeit less extensive, network changes suggesting that amnesic MCJ might be indeed an intermediate stage between healthy aging and dementia. These findings agree with the results obtained in previous studies [7, 17].

The assessment of the nodal degree in AD showed widespread changes in the number of connections of regions that belong to the default-mode network, including the medial orbitofrontal, the anterior cingulate and posterior cingulate gyri, compared to controls or patients with MCI. This network has been strongly associated with AD as its regions coincide with the areas showing amyloid deposition, gray matter atrophy and glucose hypometabolism in these patients [33]. Hence, the changes we found in this study may partially reflect pathological and metabolic abnormalities that usually occur in AD patients.

In contrast to the nodal degree, the nodal local efficiency showed alterations both in MCI and AD patients. Whereas in MCI patients, these changes were confined to a single region in the left temporal lobe, in AD patients the local efficiency was altered across several frontal, temporal and parietal areas, including the hippocampus and amygdala, which are involved in AD pathology [34]. The local efficiency reflects how efficiently is the communication between a region and its neighboring areas. Decreases in this measure might indicate a loss of local connections, whereas increases could reflect a compensatory mechanism by which the number of connections between close brain areas increases to compensate for the loss of connections between distant brain areas. Hence, altogether our findings indicate that graph theory is a useful method to assess abnormalities in brain connectivity and topology in the prodromal and clinical stages of AD.

### 4.2. Large-scale functional networks in PD-MCI

Cognitive impairment is one of the most important non-motor symptoms in PD that greatly affects quality of life. During the course of the disease, most PD patients will develop impairment in one or more cognitive domains, for which they will receive a diagnosis of MCI. The presence of MCI in PD is associated with an increased risk to progress to dementia [35]. Hence, there is a pressing need to identify the underlying mechanisms of MCI to prevent cognitive decline in PD patients. Using a weighted network approach in BRAPH, we found that PD-MCI patients presented a lower number of connections in the whole brain network compared to controls. This indicates that in general their regions were more disconnected. However, after identifying the modular structure and performing the same analysis within each module, we observed that these effects were mostly driven by a lower degree in the fronto-parietal network in the PD-MCI group. The regions that were most affected in this network were the superior frontal gyri, superior parietal gyri and precuneus. All of these regions have been previously shown to display reduced connectivity in PD [8]. In addition, they have also been identified as important brain hubs in previous graph theory studies [27]. Within the graph theory framework, the brain hubs are the most important and central regions of a network as they mediate numerous long-distance connections. This characteristic also suggests they might have higher metabolic costs and a greater vulnerability to oxidative stress [27]. In a previous study [36], it was shown that the pathological brain lesions are concentrated in hub regions of the connectome in several neurodegenerative disorders, including PD, in line with our findings.

### 4.3. BRAPH features

The application of graph theory to the field of imaging connectomics is still in its early beginnings. There are several important challenges that need to be addressed such as whether the nodes and edges are an accurate description of the true underlying brain connectome in all its complexity of millions of neurons and synapses. Although we have limited knowledge on how to address this particular issue, there are several other challenges that can be addressed through the use of BRAPH. For instance, given that the true connectome is a sparse network, it is important to threshold the structural or functional edges, which typically consist of continuous association indices [37]. This thresholding can be carried out using different methods. On the one hand, a threshold can be applied so that only the connections that are below a significance level are included in the analysis. In this way, the weaker connections of the graph are eliminated and considered spurious. This approach will yield different numbers of connections across different individuals or groups. On the other hand, a threshold can be applied such that all networks have the same number of connections through a fixed value of density. In this way, only a percentage of edges are included in the analyses and the graph theory measures are independent of the number of edges [37]. In BRAPH, both thresholding options are available so that the user can easily compare them.

Another important challenge is the choice of a given value of threshold since there is currently no way to establish which is the best value. To solve this issue, BRAPH allows testing a hypothesis across different levels of significance or densities to determine the robustness of the results. Some authors consider choosing a range of thresholds an arbitrary process that produces values that are strongly dependent on this choice. For this reason, we also provide an option for weighted network analysis, which allows assessing both strong and weak connections present in a graph and is not dependent on a particular thresholding scheme.

Another challenge that is beginning to emerge in the scientific community is the realization that different node definitions can lead to different network findings. Currently BRAPH provides six anatomical and functional brain atlases that the user can apply within the same study to assess the consistency of the results against different parcellation schemes. In addition, a new brain atlas can be easily created or uploaded in BRAPH that adjusts to the user needs.

Finally, the last challenge that can be addressed through BRAPH is the normalization of graph theory measures by reference to random networks that have a random organization (with the same degree and/or weight distribution). BRAPH provides various options to perform this normalization.

In conclusion, the study of the brain connectome is a growing field that will provide important insights into brain organization in health and disease. Amongst its numerous applications, there is the possibility it might help predicting the pathological spread of disease proteins in neurodegenerative disorders, which are becoming increasingly prevalent in the world’s aging population. To address the increasing demands in this growing field we provide BRAPH, the first object-oriented software that will integrate new topological analyses and measures in a structured manner, allowing the users to be updated with the latest developments in graph theory. BRAPH can be found at https://www.braph.org with online videos, a comprehensive manual, a support forum and relevant links. It is free for all researchers and can be used in all operating systems.

## 5. Acknowledgements

Data used in the preparation of this article were obtained from the Alzheimer’s Disease Neuroimaging Initiative (ADNI) and the Parkinson’s Progression Markers Initiative (PPMI).

Data collection and sharing of ADNI was funded by the National Institutes of Health Grant U01 AG024904 and Department of Defense award number W81XWH-12-2-0012. ADNI is funded by the National Institute on Aging, the National Institute of Biomedical Imaging and Bioengineering, and through generous contributions from the following: Alzheimer’s Association; Alzheimer’s Drug Discovery Foundation; BioClinica, Inc.; Biogen Idec Inc.; Bristol-Myers Squibb Company; Eisai Inc.; Elan Pharmaceuticals, Inc.; Eli Lilly and Company; F. Hoffmann-La Roche Ltd and its affiliated company Genentech, Inc.; GE Healthcare; Innogenetics, N. V.; IXICO Ltd.; Janssen Alzheimer Immunotherapy Research & Development, LLC.; Johnson & Johnson Pharmaceutical Research & Development LLC.; Medpace, Inc.; Merck & Co., Inc.; Meso Scale Diagnostics, LLC.; NeuroRx Research; Novartis Pharmaceuticals Corporation; Pfizer Inc.; Piramal Imaging; Servier; Synarc Inc.; and Takeda Pharmaceutical Company. The Canadian Institutes of Health Research is providing funds to support ADNI clinical sites in Canada. Private sector contributions are facilitated by the Foundation for the National Institutes of Health (http://www.fnih.org). The grantee organization is the Northern California Institute for Research and Education, and the study is coordinated by the Alzheimer’s Disease Cooperative Study at the University of California, San Diego. ADNI data are disseminated by the Laboratory for Neuro Imaging at the University of California, Los Angeles.

PPMI - a public-private partnership - is funded by the Michael J. Fox Foundation for Parkinson’s Research and funding partners, including Abbott, Avid Radiopharmaceuticals, Biogen Idec, Bristol-Myers Squibb, Covance, Elan, GE Healthcare, Genentech, GSK-GlaxoSmithKline, Lilly, Merck, MSD-Meso Scale Discovery, Pfizer, Roche, UCB (http://www.ppmi-info.org/fundingpartners). For up-to-date information on the PPMI database visit http://www.ppmi-info.org.

We would also like to thank the Swedish Foundation for Strategic Research (SSF), the Strategic Research Programme in Neuroscience at Karolinska Institutet (StratNeuro), Hjärnfonden, and Birgitta och Sten Westerberg for additional financial support.

